## Extended Data Figure 1 for "Targeting properdin - Structure and function of a novel family of tick-derived complement inhibitors"

a) MonoQ fractionation

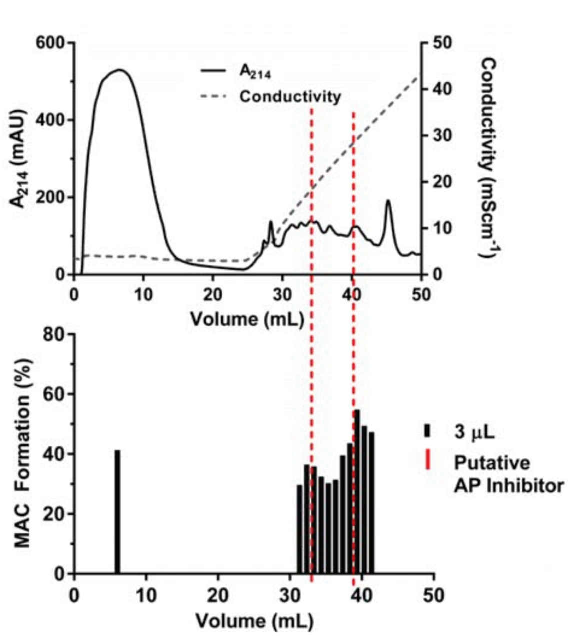


b) SEC fractionation

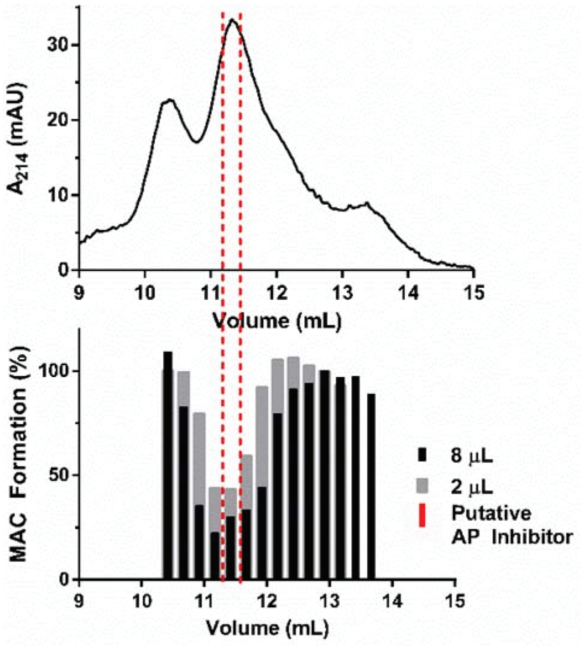


c)

| Accession No. | Re-name | Score | Mature Protein Mass (Da) | pI | BLAST Top Hit | BLAST E score | BLAST RefSeq |
|---|---|---|---|---|---|---|---|
| comp52_seq0 | AP1 | 239 | 20068 | 4.67 | None | | |
| comp1106_seq1 | AP2 | 79 | 19433 | 5.97 | Salivary lipocalin (A. variegatum) | 1.00E-13 | DAA34698.1 |
| comp3626_seq0 | AP3 | 130 | 45968 | 5.16 | Hypothetical Protein, IscW, Ixodes scapularis | 0 | XP_002409462.1 |
| comp1215_seq0 | AP4 | 110 | 38706 | 6.15 | Proliferation-associated Protein 2G4 | 1.00E-157 | KDR23244.1 |
| comp8435_seq1 | AP5 | 81 | 22430 | 4.89 | None | | |
| comp5629_seq0 | AP6 | 71 | 24630 | 8.79 | None | | |
| comp4_seq0 | AP7 | 36 | 21995 | 5.38 | Lipocalin | 5.00E-4 | ABI52661.1 |
| RpIx75-921729 | - | 612 | 22957 | 5.21 | Salivary lipocalin (A. variegatum) | 3.00E-11 | DAA34698.1 |
| RpSigP-58530 | - | 24 | 21995 | 5.38 | N(G),N(G)-di-methylarginine dimethylamino-hydrolase | 3 | WP_011213937.1 |

d)

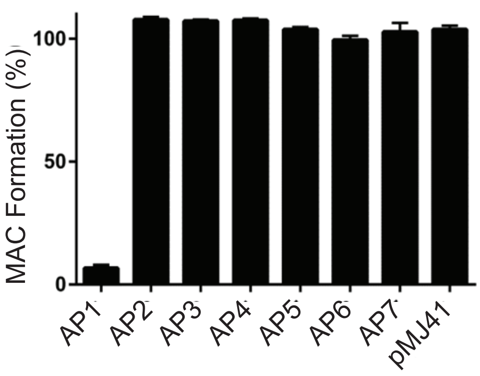
