## Extended Data Figure 2 for "Targeting properdin - Structure and function of a novel family of tick-derived complement inhibitors"

a)

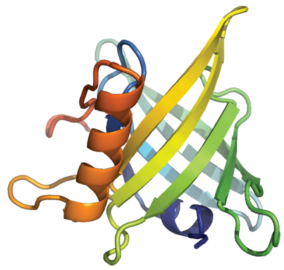

CirpA1

b)

| | CirpA1 | CirpA2 | CirpA3 | CirpA4 | CirpA5 | CirpA6 | Organism of origin |
|---|---|---|---|---|---|---|---|
| CirpA1 | 100 | 82 | 52 | 48 | 46 | 43 | *Rhipicephalus pulchellus* |
| CirpA2 | 82 | 100 | 72 | 57 | 48 | 44 | *Rhipicephalus pulchellus (m, IH)* |
| CirpA3 | 52 | 72 | 100 | 56 | 42 | 47 | *Rhipicephalus pulchellus (f, IH)* |
| CirpA4 | 48 | 57 | 56 | 100 | 42 | 44 | *Rhipicephalus appendiculatus* |
| CirpA5 | 46 | 48 | 42 | 42 | 100 | 44 | *Rhipicephalus appendiculatus (IH)* |
| CirpA6 | 43 | 44 | 47 | 44 | 44 | 100 | *Rhipicephalus microplus* |

c)

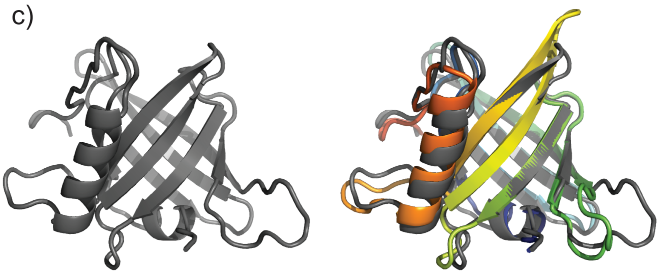

CirpA3

RMSD=0.95 Å
52% seq. identity

d)

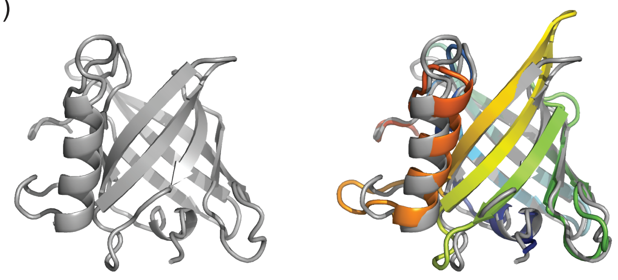

CirpA4

RMSD=0.69 Å
48% seq. identity

e)

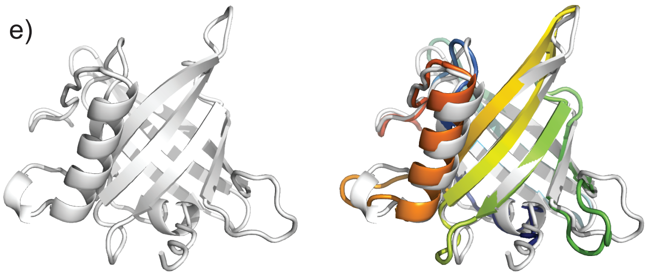

CirpA5

RMSD=0.85 Å
46% seq. identity

f)

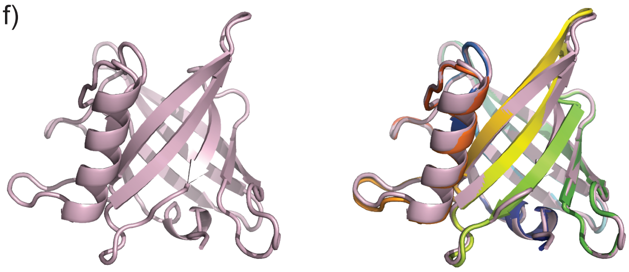

CirpA1 (complex)

RMSD=0.347 Å
